# STRIDE: Species Tree Root Inference from Gene Duplication Events

**DOI:** 10.1101/140020

**Authors:** David M. Emms, Steven Kelly

## Abstract

The correct interpretation of a phylogenetic tree is dependent on it being correctly rooted. A gene duplication event at the base of a clade of species is synapamorphic, and thus excludes the root of the species tree from that clade. We present STRIDE, a fast, effective, and outgroup-free method for species tree root inference from gene duplication events. STRIDE identifies sets of well-supported gene duplication events from cohorts of gene trees, and analyses these events to infer a probability distribution over an unrooted species tree for the location of the true root. We show that STRIDE infers the correct root of the species tree for a large range of simulated and real species sets. We demonstrate that the novel probability model implemented in STRIDE can accurately represent the ambiguity in species tree root assignment for datasets where information is limited. Furthermore, application of STRIDE to inference of the origin of the eukaryotic tree resulted in a root probability distribution that was consistent with, but unable to distinguish between, leading hypotheses for the origin of the eukaryotes. In summary, STRIDE is a fast, scalable, and effective method for species tree root inference from genome scale data.

## Introduction

*“Nothing in biology makes sense except in the light of evolution”* (Dobzhansky, T. 2013), *“nothing in evolution makes sense except in the light of phylogeny”* (Sytsma, K.J. and Pires, J.C. 2001), and nothing in a phylogeny makes sense except in the light of its root. For example, the phylogeny for four species (Fig. 1A) has five possible roots (Fig. 1B-F) and each of the different roots corresponds to a different hypothesis as to the evolutionary history of the species. For the presented tree, identifying a wrong branch as the root (for example Fig. 1E) would lead us to conclude that elephants are more closely related to fish and birds than they are to wolves, even though we are using a tree with the correct topology. A species tree only gives the correct evolutionary relationships when rooted correctly (Fig. 1B). Thus it is of critical importance to our interpretation of relationships, and the evolutionary history of life on earth, that we have accurate methods of inferring the root of species phylogenies.

**Figure 1.**
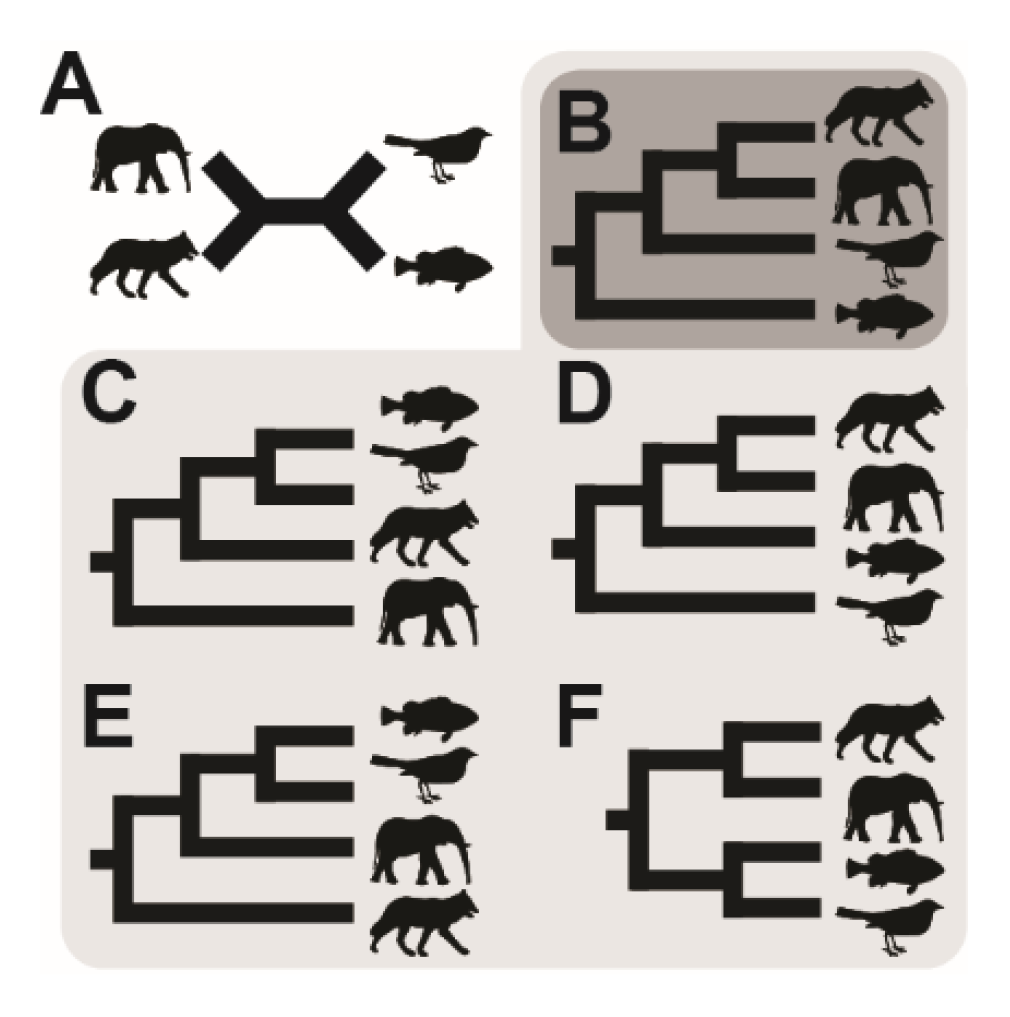
Possible roots for a four-taxa species tree. A) Unrooted species tree for four species: elephant, wolf, fish & bird. B) The correct rooting of the species tree. B-F) The five possible rooted species trees for the unrooted species tree in A.

Correct species tree rooting is also of critical importance for the inference of orthology relationships between genes. Given an unrooted gene tree (Fig. 2A), knowledge of the correct branching order of the species tree (Fig. 1B) is required to correctly root the gene tree (Fig. 2B). An incorrect rooting of the species tree (Fig. 1C-F) leads to an incorrect inference of the root of the gene tree (Fig. 2C-F), and thus incorrect identification of orthologous genes (Fig. 2G-H). Therefore, our ability to compare the biology of species, through comparisons between orthologous genes, is reliant on accurate methods of inferring the root of species phylogenies.

**Figure 2.**
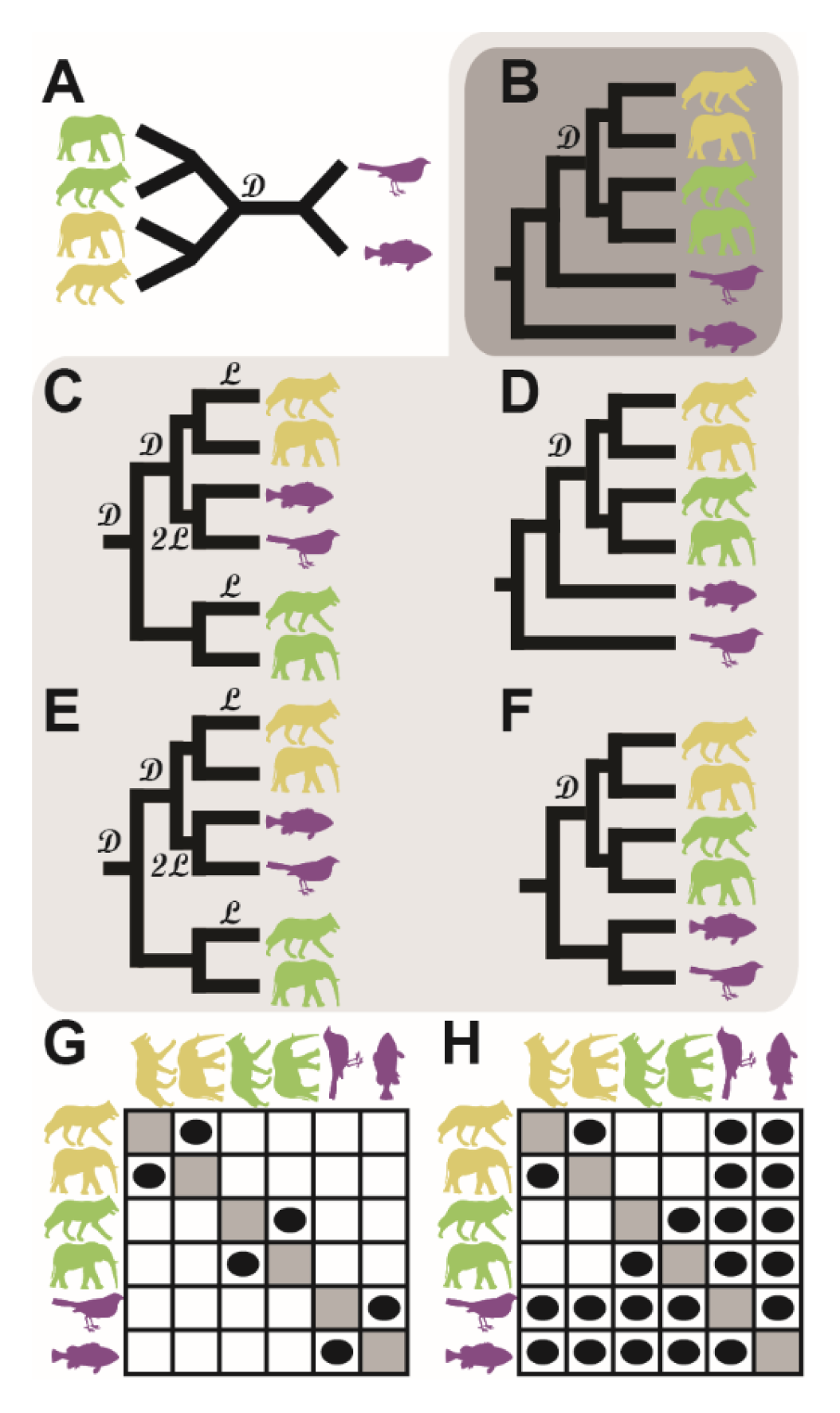
Orthologues inferred from gene trees depend on the root. A) An unrooted gene tree corresponding to an orthogroup with a gene duplication event in the common ancestor of wolf and elephant. Genes from each species are represented by an image of the species. B-F) The most parsimonious rootings of the gene trees (fewest duplications and losses) for each of the five roots of the species tree, as shown in Figure 1 B-F. 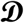 - gene duplication event, 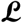 - gene loss event. G) Orthologues inferred from the incorrect trees D & E. H) Orthologues inferred from the correctly rooted tree B and also the close to correctly rooted trees D & F.

Although correct root placement is essential for our ability to interpret phylogenies, almost all models of sequence evolution used for tree inference are time-reversible and produce unrooted phylogenetic trees. In order to identify the root of a phylogeny extra information is required, usually knowledge of an extra species that is a suitable (i.e. closely related) outgroup for the set of species for which the root is unknown. However, outgroup choice is a common source of error in phylogenetic tree inference, with distantly related outgroups leading to inaccurate root placement and distortion of the phylogeny due to long branch attraction (Felsenstein, J. 1981) (Berger, S.A., Krompass, D., et al. 2011). While time-irreversible models of sequence evolution exist, they do not implicitly provide a method for accurately inferring the direction of time in a tree (Huelsenbeck, J.P., Bollback, J.P., et al. 2002, Williams, T.A., Heaps, S.E., et al. 2015). To address this issue, methods have been developed that can simultaneously infer rooted species and gene trees (Boussau, B., Szollosi, G.J., et al. 2013). However, these methods are computationally expensive and do not scale well to moderate or large species sets.

“Duplicate gene rooting” has been proposed as an alternative method for rooting species trees (Donoghue, M.J. and Mathews, S. 1998, Simmons, M.P., Bailey, C.D., et al. 2000). The conceptual basis for this approach is that gene duplication events are time-irreversible, unlike character substitution, and thus infer the direction of time on the species tree. Specifically, every node in an unrooted, binary gene tree has three branches incident upon it. If the node is a speciation node then any of the three incident branches could be the edge in the direction of the root, with the other two being in the opposite direction. Thus, speciation nodes are uninformative about the direction of time along the tree. For a duplication node, however, the symmetry is broken. Two of the edges will correspond to the two copies of the gene post-duplication, while the third edge will correspond to the gene pre-duplication and thus point towards the root of the tree (Fig. 2A, node marked 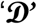). In the case of this example tree, it can be inferred that the root of the species tree must be outside of the subtree containing elephant and wolf. In an idealised case (with no effects such as incomplete lineage sorting or lateral gene transfer) the two post-duplication branches can be distinguished from the pre-duplication branch as the post-duplication branches contain genes from overlapping species sets. Furthermore, these species sets will be identical if there has been no gene loss or horizontal gene transfer, and the topology of the duplicate subtrees will recapitulate the species tree topology. Thus, if gene duplication nodes can be accurately identified in an unrooted gene tree, then the direction of time can be ascertained for all branches in the post-duplication subtrees. The direction of time on these branches determines the direction of time on the corresponding branches of the species tree, and multiple gene duplication events can be aggregated to determine the direction of time across the whole species tree, thus revealing the location of the root.

Here we present STRIDE, a novel algorithm for <u>S</u>pecies <u>T</u>ree <u>R</u>oot <u>I</u>nference from gene <u>D</u>uplication <u>E</u>vents. STRIDE identifies sets of well-supported gene duplication events from cohorts of gene trees, and analyses these events to infer a probability distribution over an unrooted species tree for the location of the true root. We show that STRIDE correctly identifies the community-accepted root of the majority of species trees. Additionally, we demonstrate that STRIDE effectively captures uncertainty in root placement when data is limited or conflicting. Finally, we demonstrate the utility of STRIDE to challenging phylogenetic problems by providing an outgroup-free root analysis of the origin of the eukaryotes.

## Methods

### Problem definition and approach

A branch of an unrooted species tree corresponds to a bipartition that splits the tree’s taxa into two blocks. The presence of a well-supported gene duplication that respects the topology of the species tree is a synapamorphy that stipulates that the block in which the duplicates are found is a monophyletic clade. This synapamorphy identifies the direction of time along the branches within this monophyletic clade. The single exception to this is the branch in the unrooted tree corresponding to the root in which time flows in both directions. This is because the branch that spans the root in the unrooted species tree corresponds to two branches in the rooted species tree and both of its corresponding blocks are monophyletic clades (Fig. 1A & B). The method presented here aims to identify this root branch by identifying and analysing a set of well-supported gene-duplication events. The method identifies the complete set of gene duplication events contained within a set of user-supplied gene trees and uses these to infer the location of the root of the species tree. To express uncertainty in the case of limited data or data conflict, the method uses a probabilistic model of gene-duplication events to calculate a probability distribution across the branches of the species tree for the location of the root.

### Inference of Orthogroups and Gene Trees

For each species set, the protein sequence translations of representative gene models were downloaded from appropriate online databases. These protein sequences were then subject to orthogroup inference using OrthoFinder v1.1.4 (Emms, D.M. and Kelly, S. 2015). The resulting sets of protein sequence orthogroups were aligned using MAFFT L-INS-I v7.305b (Katoh, K. and Standley, D.M. 2013) and subject phylogenetic inference using IQTREE v1.5.3 (Nguyen, L.T., Schmidt, H.A., et al. 2015). All methods used their default settings. Parallelisation of MAFFT and IQTREE runs was done using GNU Parallel (Tange, O. 2011). Alignments were viewed using AliView (Larsson, A. 2014). Trees were viewed using Dendroscope (Huson, D.H. and Scornavacca, C. 2012) and drawn using the ETE library (Huerta-Cepas, J., Serra, F., et al. 2016).

### Identification of Putative Duplications

Gene-duplication events are considered informative if they occur on any branch other than a terminal branch in the species phylogeny, as a duplicated gene that occurs only in a single species is not informative to the position of the root of the tree. To identify informative gene duplication events a novel tree analysis algorithm was developed (Fig. 3). Prior to analysis of the gene trees, the unrooted species tree was analysed to determine the species sets in which genes would be expected to occur following a gene duplication event along any branch of the species tree. For each direction along each branch the sets of species in the child clades (X & Y) and in the grandchild clades (x_1_, x_2_, y_1_ & y_2_) immediately following that branch were identified (Fig. 3A). Then each gene tree was analysed in turn to identify well-supported gene duplication events within the gene tree as follows: each node, n, in the tree was considered in turn, if the node was an unresolved polytomy it was excluded as such nodes correspond to either a higher-order multiplication events (e.g. triplication) or an unresolved event in the gene tree (e.g. an amalgamation of several weakly supported bipartitions). Each analysed node therefore had three branches incident on it, and any pair of branches could potentially represent duplicate genes (Fig. 3B). For each pair of branches, b_1_ and b_2_, the sets of species, S_1_ and S_2_, below each branch were used to identify the locations in the species tree corresponding to these branches in the gene tree. This was done by identifying the smallest block, B_i_ in the species tree that contains all the species in S_i_ (i=1,2), thus making the method robust in the case of subsequent gene loss (Fig. 3B). If there was more than one block satisfying this criteria, each of these possible blocks were considered. A node, n, was considered as a putative gene duplication event if B_1_=B_2_.

**Figure 3.**
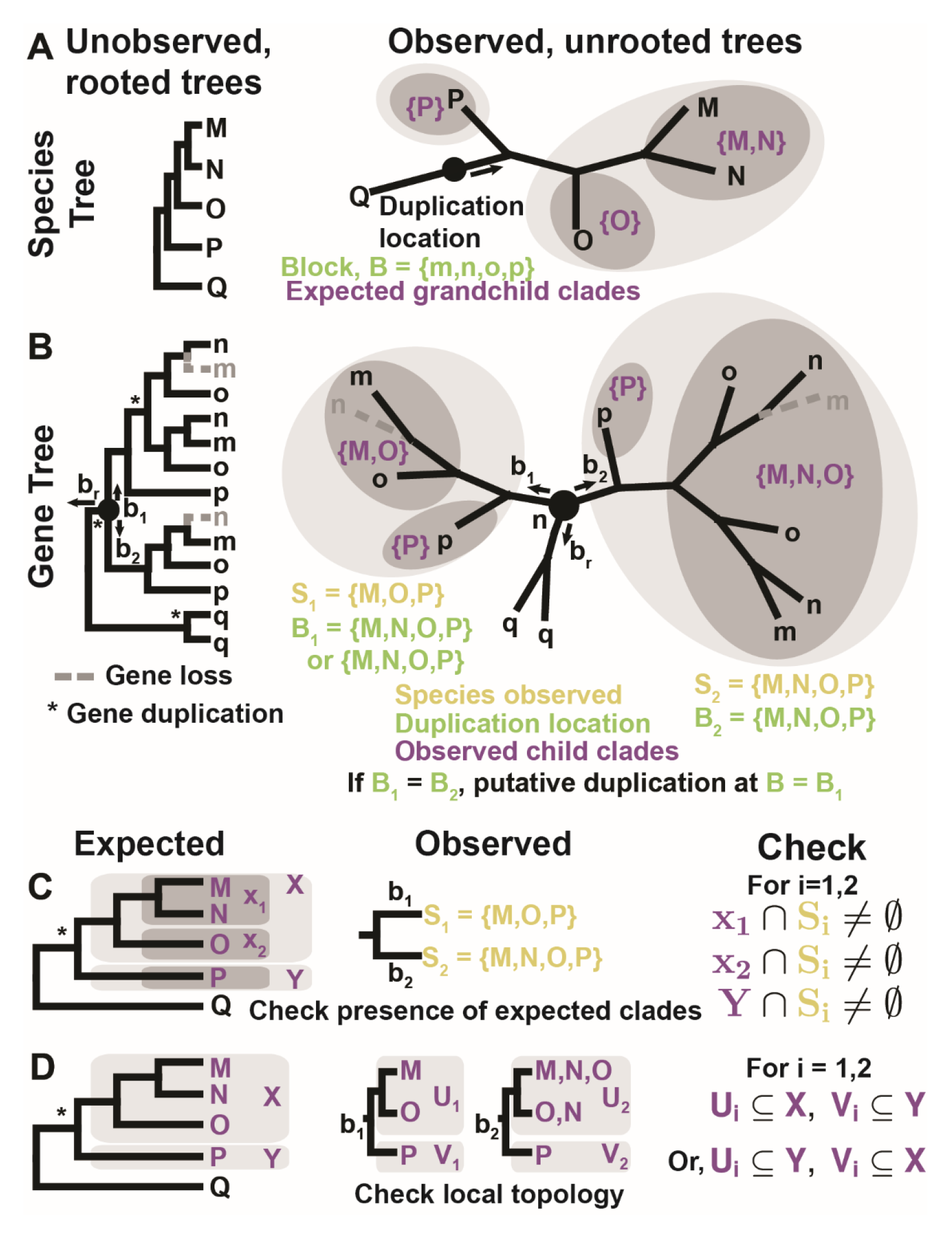
Identification of well-supported gene duplication events. Upper case letters M,N,O,P & Q are species, lower case m,n,o,p & q are genes from the corresponding species. A) The unknown, rooted species tree (left) and the observed, unrooted species tree (right). Black dot and arrow show the location of a single hypothetical gene duplication event on a branch with time flowing in the direction indicated by the arrow. The branch location and direction is uniquely identified by the block, B, of species whose common ancestor would have inherited the duplicate genes. The expected species in the child clades (X & Y) and grandchild clades (x_1_, x_2_ & Y) after this hypothetical duplication are highlighted with light/dark grey ellipses respectively. B) The unknown, rooted gene tree (left) and the observed, unrooted gene tree (right) for a hypothetical gene family with three gene duplication events (marked by *) and two gene loss events (grey, dotted line). The node currently being analysed is n and br is the current, tentative direction towards the root. For these n and b_r_, b_1_ and b_2_ are analysed to see if they are well-supported gene duplication branches. Si is the set of species below branch b_i_, B_i_ is the smallest block of a bipartition containing S_i_ (i=1,2) C) The check that genes from each of the expected grandchild clades are present on each duplicate branch D) The check that the local topology for each duplication branch agrees with expected topology. U_i_ and V_i_ are the observed child clades on branch b_i_. The observed child clades should not contain genes from any species not in the expected child clades

Nodes that were identified as putative gene duplication events were further examined to reduce the possibility that their existence or location had been misidentified due to errors in gene tree inference. The criteria were: 1) There must be at least one gene from each of the expected grandchild clades in both S_1_ and S_2_ (Fig. 3C). 2) The branching structure immediately after the gene duplication event on branches b_1_ and b_2_ must match the expected branching structure (Fig. 3D), i.e. the first node for each duplicate split the descendent species into the expected sets X and Y, or subsets thereof. Note that it would not be valid to check the topology to the level of grandchild clades in step 2 since this would fail to identify gene duplication events if there were also a subsequent gene duplication event one branch lower in the species tree. In this case, the observed grandchild clades would both be subsets of one of the expected child clades rather than grandchild clades. Steps 1 and 2 check that the observed clades are subsets of the expected clades (rather than requiring they be equal to) as this is necessary to make the method robust to subsequent gene loss events.

### Identifying the Maximum-Parsimony Root of the Species Tree

As discussed above, a gene duplication on a bipartition of an unrooted species tree i stipulates the direction of time for all branches of the subtrees derived from that bipartition. Given a set of gene duplication events, the branch in the species tree that violates the fewest gene duplication events is identified as the maximum parsimony root. If multiple such branches exist then they are each identified as equally parsimonious.

### Probability model for the root of the species tree

For any given set of gene-trees, it is possible that errors in gene-tree inference will lead to false positive inference of gene duplication events that past the filtration criteria. To account for this, a probability model was developed for the location of the root of the tree given the set of (potentially conflicting) putative gene duplication events identified. The model consisted of two parts. The first part, the branch-level model, calculated the probability that a branch was the root given only the duplications observed in either direction along that branch. The second part, the tree-level model, aggregated all duplications observed across all branches of tree to give the final probability distribution for the location of the root taking into account all information obtained from all gene duplication events observed across the tree.

At the branch-level, the set of putative gene duplication events identified on that branch are modelled by two Poisson processes, one giving rise to true positive gene duplications and the other to false positive duplications. On a given branch, *i*, of a species tree, *m_i_* duplications are observed that support time flowing in one direction along the branch, →, and *n_i_* duplications supporting time flowing in the opposite direction, ←. The set of duplications on branch *i* is then written, 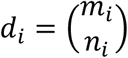, and *D* is the set of putative duplications observed on all branches of the species tree, *D* = [*d*_1_, *d*_2_,…, *d_b_*}. The final tree-level probability distribution *P(i = root\D)* takes into account the complete set of duplications, D, observed on all branches of the tree rather than just the duplications, di, observed on a single branch:

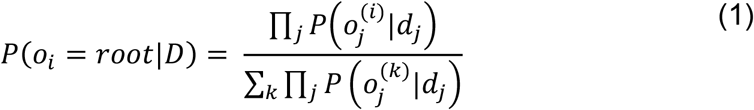

where 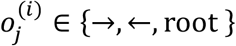 is the orientation of the branch *j* that would be implied by the root of the tree being branch *i*. That is, the probability distribution for the root given all the gene duplication events on the tree can be expressed in terms of the probabilities for the orientation of each branch given only the gene duplications on that branch; *P*(→ |*d_i_*), *P*(← |*d_i_*) and *P*(root|*d_i_*).

### Poisson Model for Gene Duplications

To calculate *P*(*o_i_*|*d_i_*) the duplications observed on a branch are modelled as arising from two Poisson processes. One process describes the number of true positive duplications (corresponding to the actual direction of time along the branch) and the other describes the number of false positive duplications. Let α be a parameter giving the relative frequency of false positives to true positives across all branches of the tree. Then *m*~*P_o_*(*λ*) and *n*~*P_o_*(*αλ*), where λ is the expected number of true positives on the branch. We set the total expected number of duplications on the branch from the two Poisson processes to match the actual number observed, N. Thus *λ* = *N*/( 1 + *α*). The relative rate of false positives to true positives across the whole tree can be estimated from the number conflicting duplications given the maximum parsimony root of the tree. So as not to over-penalise false-positive duplications, we take α to be one tenth of the ratio of conflicting to non-conflicting duplications of the maximum parsimony root.

Bayes’ rule gives

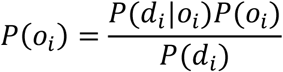

where 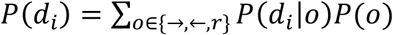. The priors are given by 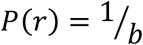 and 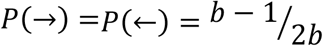, where b=2t-3 is the number of branches on an unrooted tree with t taxa. The probability mass function for the Poisson distribution immediately gives *P*(*d*|→) and *P*(*d*|←:

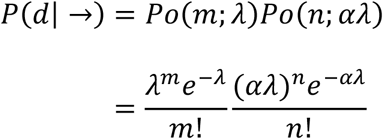

and,

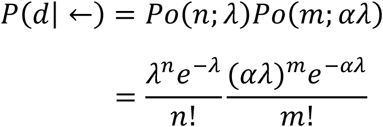

The branch with the root is more complicated since it actually corresponds to two branches on the rooted tree we are attempting to recover. On these two branches time flows in opposite directions, away from a central root that separates them. We must allow for the 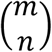 duplications on the branch to actually correspond to 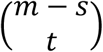 duplications on one of the two branches and 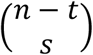 on the other branch (with opposite orientation to the first). The number of false positive duplications, *s* and *t*, are unknown and therefore must be summed over. Similarly, the location of root could fall at any point along the length of the original branch. If the root were a fraction, *x*, along the length of the branch then the expected rate of false positive and true positive duplications on that fraction of the branch would be *xλ* and *xαλ* respectively whereas on the other branch the rates would be *(1-x)λ* and *(1-x)αλ*. Thus, integrating over the position of the root along the branch and summing over the distribution of the 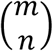 putative duplications between true positives and false positives on the two resulting branches, we find:

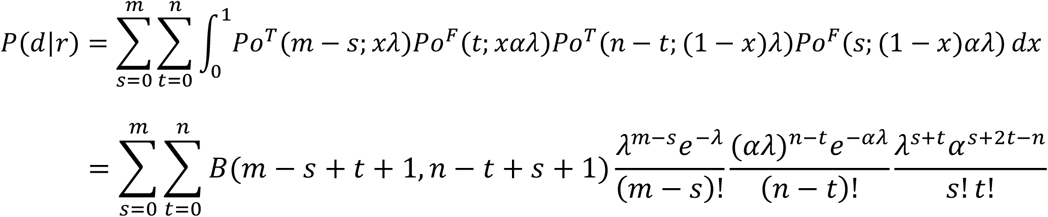

Where *B(,)* is the beta function.

The duplications observed in just one species are uninformative as to the location of the root and so should not affect the root probabilities produced by the model. As such, the branch model for terminal branches is modified to only model the number of inward duplications (those supporting the tree minus the species on the terminal branch as a monophyletic clade). The rates *λ_Term,TP_* and *λ_Term,FP_* are the observed true positive and false positive rates for inward duplications on the terminal branches for the maximum parsimony root. For the terminal branches, the branch model is:

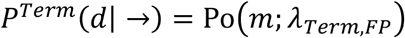

and

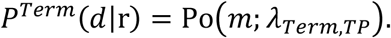

The branch-level model takes into account only the duplications observed on a single branch and these probabilities feed into the tree-level model to give the final probabilities for the position of the root (Fig. 4). The behaviour of the branch model is in good agreement with an intuitive understanding of the probabilities that should be assigned to the three possible orientations for a branch given the number of putative duplications observed in either direction (Fig. 4A-C). The probability of time flowing to the left/right increases monotonically with the number of putative duplications supporting it. The probability of a branch being a root is highest when the number of putative gene duplications in both directions is the same. Finally, the probability of a branch being a root remains significantly above zero if there is any number of gene duplications in both directions (Fig. 4B & C). This reflects the fact that putative gene duplications supporting the monophyletic nature of both blocks of a bipartition support that bipartition being the root. The fact that there could be a large difference in the number of gene duplications in one direction compared to the other due to different branch lengths on the two sides of the root is accounted for by integrating over the position of the root along the original root branch. Thus, the probability of a branch being a root is > 30% when there are 20 duplications in one direction compared to 5 in the opposite direction (Fig. 4C). For comparison, the probability of the orientation of the branch being in the direction of the 5 duplications is vanishingly small (~10^−13^). The branch-level probability model thus gives probabilities for each branch taking into account only the duplications observed on that branch. The final probabilities for the root of the tree, taking into account all duplications across the tree are then given by the tree-level model (Equation 1, Fig. 4D & E).

**Figure 4.**
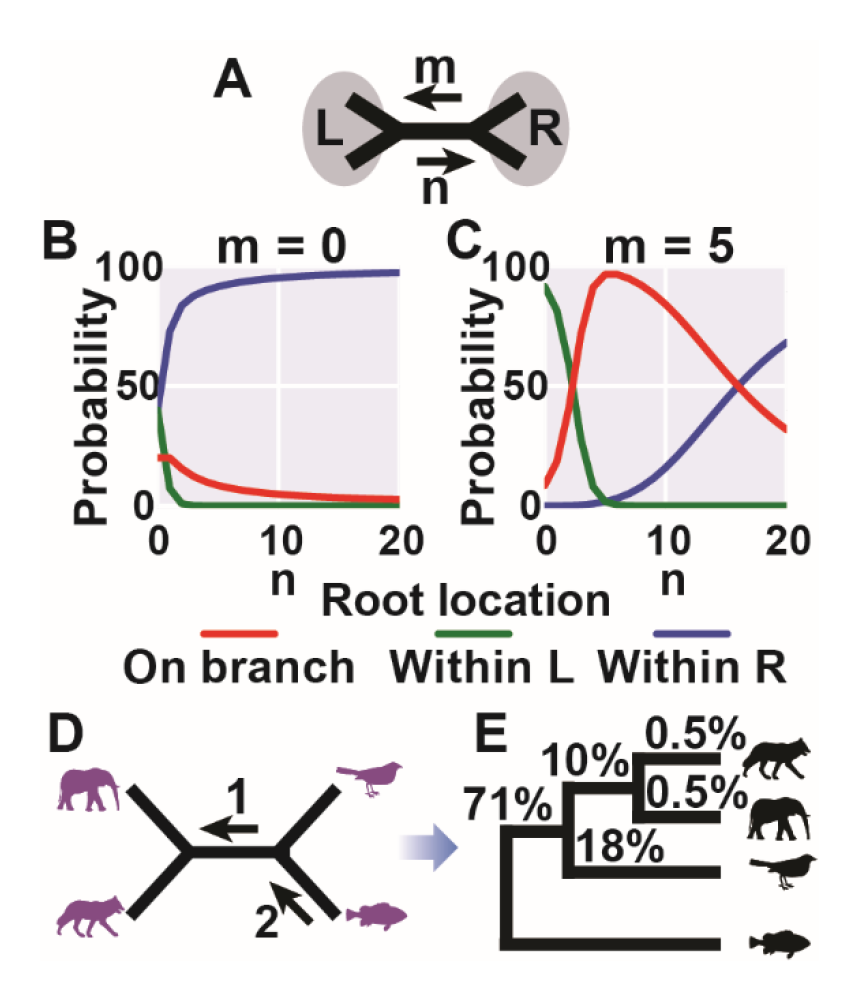
The branch-level probability model employed by STRIDE. These branch-level probabilities are used by the tree probability model to give the overall probabilities for the location of the root of the species tree. A) A single branch in the tree with m/n duplications supporting L/R as monophyletic clades. B) Branch-level model probabilities for position of the root with respect to the branch when m=0 (the model only takes into account duplications on that branch). C) As for B with m=5. D) Hypothetical total number of gene duplication events on the 4 species phylogeny. One gene duplication event is shared by elephant and dog and 2 are shared by elephant, dog and bird. D) The final tree-level model probabilities for the location of the root calculated by STRIDE taking into account all the gene duplication events on all branches in D.

### Algorithm implementation and availability

STRIDE is implemented in python. Further information, use instructions, an example dataset, and a standalone implementation of the algorithm is available under the University of Oxford Academic Use Licence at https://github.com/davidemms/STRIDE. The complete set of gene trees and species trees required to replicate this analysis are provided for download form the Zenodo research data archive at https://doi.org/10.5281/zenodo.581360.

## Results

### STRIDE identifies the correct root of species trees given simulated gene tree datasets

The ability of STRIDE to correctly infer the root of a known species tree was tested using three published, simulated gene tree datasets. The first dataset consisted of 2000 simulated gene trees from 40 species with heterogeneous rates of gene duplication and loss within trees (Boussau, B., Szollosi, G.J., et al. 2013). The second and third datasets consisted of 12000 gene trees from 12 species and 7500 gene trees from 17 species, respectively. These two datasets were similar to the first dataset but also incorporated incomplete lineage sorting generated using a range of effective population sizes (Wu, Y.C., Rasmussen, M.D., et al. 2014). Since incomplete lineage sorting can lead to misidentification of gene duplication and loss events these latter two datasets provided a good test of STRIDE’s robustness in the face of gene-tree/species-tree incongruence. For all three simulated datasets, STRIDE correctly inferred the root of the species tree and assigned it a probability of 100% (Table 1, Supplementary File 1. Fig. S1-S3). Thus for these simulated datasets the method performed well.

**Table 1.**
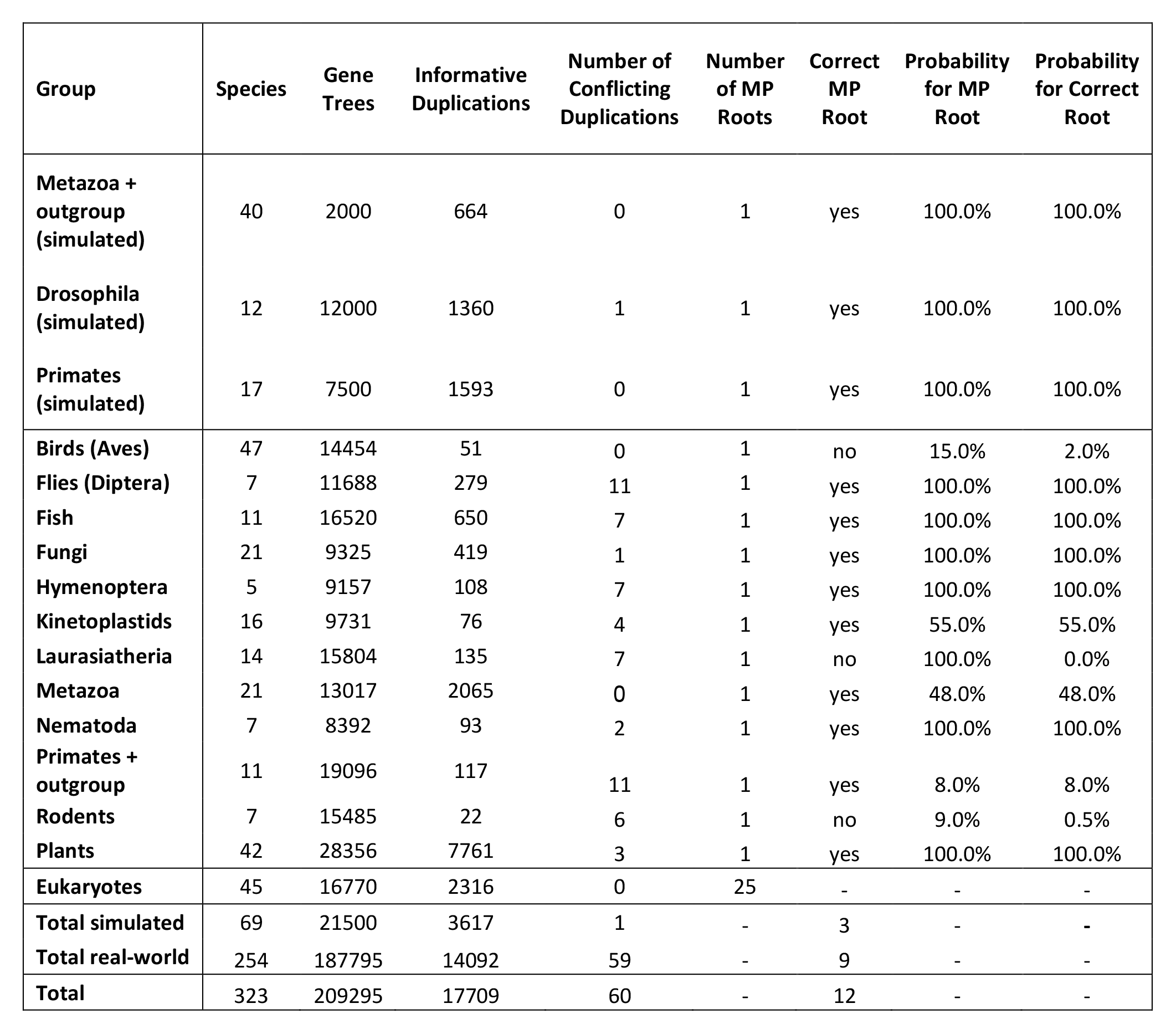

### Application of STRIDE to real species datasets

Simulated datasets generally do not capture all the nuances and difficulties seen in real biological datasets. These nuances include errors in orthogroup inference, alignment inference and gene tree inference. Thus to demonstrate the utility of STRIDE, a diverse range of groups of species were sampled from throughout the eukaryotic domain (Table 1). This included every group of eukaryotes on Ensembl Genomes containing more than 4 genera (Yates, A., Akanni, W., et al. 2016). To expand this group of tests, additional sets of genomes were obtained for 47 Birds (Jarvis, E.D., Mirarab, S., et al. 2014), 42 Green Plants (Goodstein, D.M., Shu, S.Q., et al. 2012) and 16 Kinetoplastids (Aslett, M., Aurrecoechea, C., et al. 2010). In total, this gave 12 species groups with varying levels of taxon sampling and with estimated divergence times ranging from c. 56 million years for the Primates (dos Reis, M., Donoghue, P.C.J., et al. 2014) to c. 1500 million years for the Green Plants (Parfrey, L.W., Lahr, D.J.G., et al. 2011). These species sets thus provided a diverse group with which to test the utility of STRIDE. Furthermore, for each of these species sets, there is an accepted consensus on the topology and location of the root of the species tree (Supplemental File 1). In all cases these topologies and root branches were assumed to be true when STRIDE’S performance was assessed. On average, across each of the simulated and real dataset in this analysis STRIDE took ~18 seconds to run using four cores of an Intel Core i7-4770 3.4GHz CPU.

Orthogroups for each species set were inferred using OrthoFinder (Emms, D.M. and Kelly, S. 2015), and gene trees for each orthogroup were inferred using IQTREE v1.5.3 (Nguyen, L.T., Schmidt, H.A., et al. 2015) from a multiple sequence alignment generated using MAFFT L-INS-I v7.305b (Katoh, K. and Standley, D.M. 2013). For each species set, STRIDE was run with a published unrooted species tree (without branch lengths) and the complete set of gene trees inferred from all orthogroups identified by OrthoFinder. The number species, gene trees, informative duplications and other details are provided in Table 1.

In all 12 test cases, there is a single maximum parsimony root. In 9 of the 12 tests this root agreed with the accepted root of the species set (Table 1). Figures 5 to 7 present the results of the STRIDE analysis applied to the plant, fungi, and bird datasets. These datasets correspond to the largest, median and smallest number of informative duplications per species identified by STRIDE. The results for the remaining datasets can be found in Supplemental File 1 Figures S4-S12. For the plant dataset, sufficient gene-duplication events were identified for the probability model to assign a probability of 100% to the accepted root separating the algae from the land plants (Ruhfel et al. BMC Evolutionary Biology201414:23) (Fig. 5). A probability of 100% was also assigned for the correct root in the fungi, even though fewer informative gene duplication events were identified (Fig. 6, Table 1). In both the plant and fungal datasets, STRIDE also identified substantial numbers of gene duplication events that support sub-clades within both species trees (Fig. 5 and Fig. 6).

**Figure 5.**
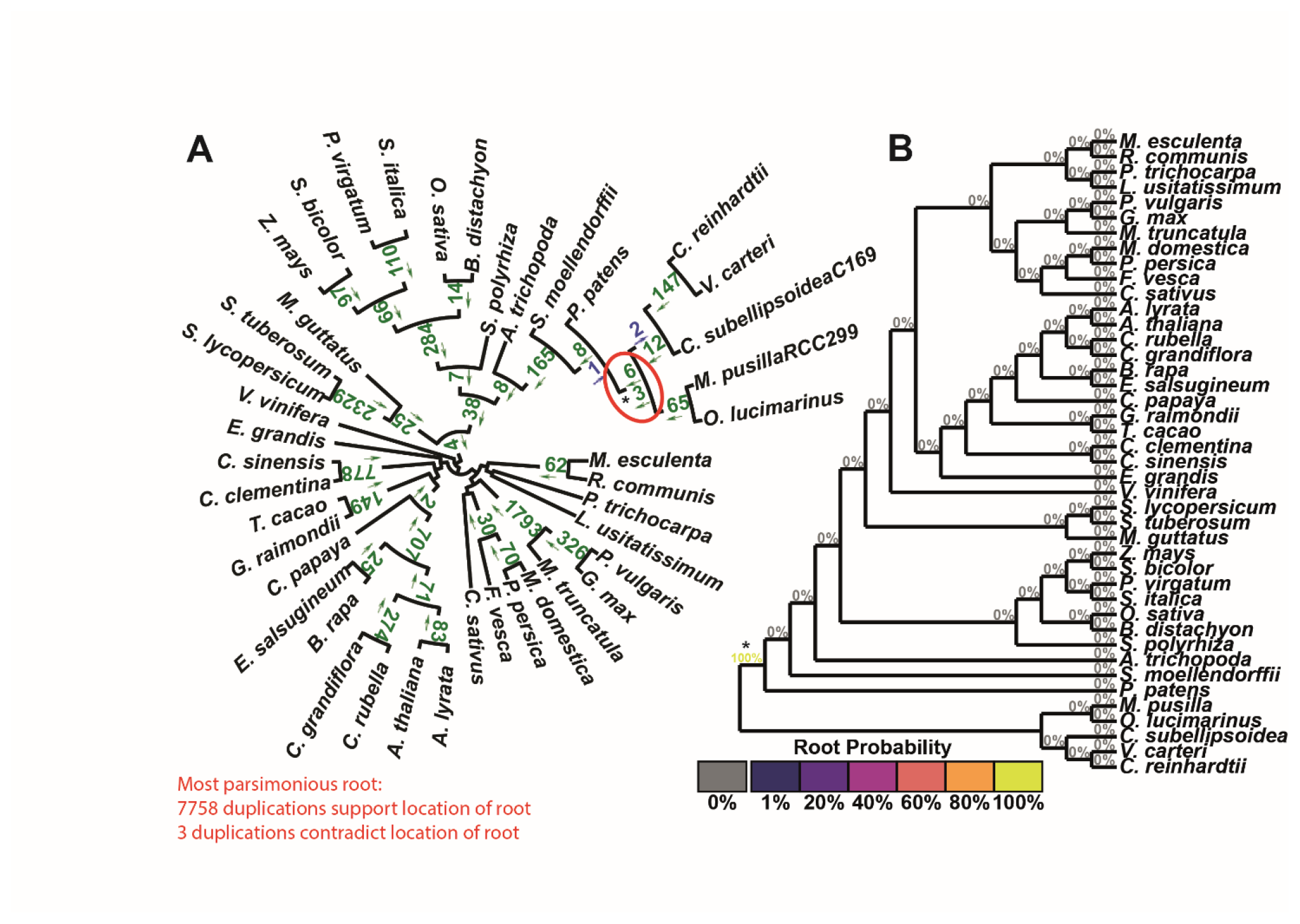
STRIDE analysis applied the set of plant gene trees. A) Numbers of identified gene duplication events are marked on the branches they are observed on and arrows indicate the direction in which the duplication occurred. Gene duplication events are in agreement with the maximum parsimony root of the tree if the arrow points away from the root, and are in green. Those that disagree are in blue. The maximum parsimony root is circled in red and is in agreement with the correct root, marked with a *. B) The probabilities for the location of the root calculated by STRIDE.

**Figure 6.**
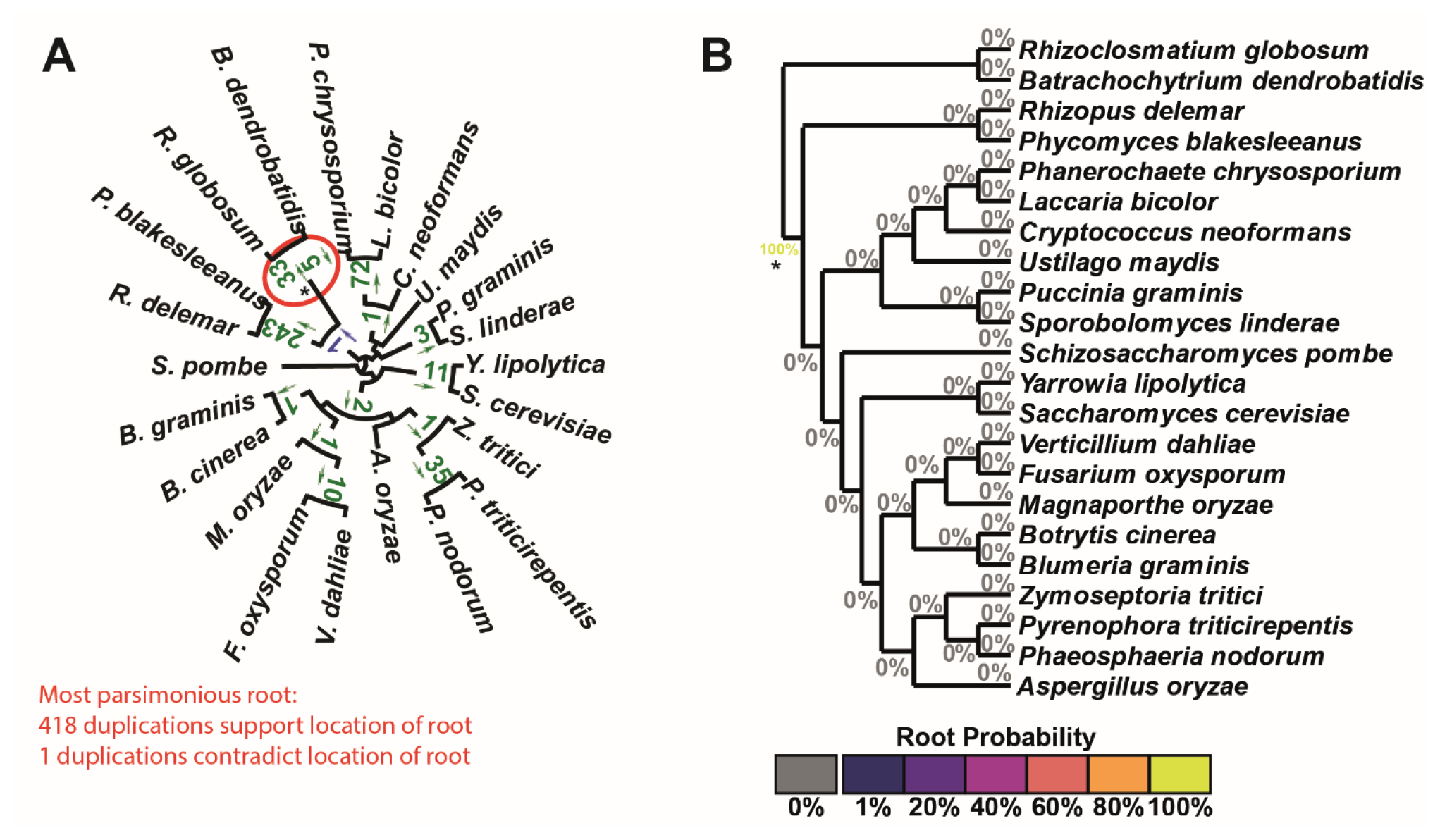
STRIDE analysis applied the set of fungi gene trees. A) Numbers of identified gene duplication events are marked on the branches they are observed on and arrows indicate the direction in which the duplication occurred. Gene duplication events are in agreement with the maximum parsimony root of the tree if the arrow points away from the root, and are in green. Those that disagree are in blue. The maximum parsimony root is circled in red and is in agreement with the correct root, marked with a *. B) The probabilities for the location of the root calculated by STRIDE.

**Figure 7.**
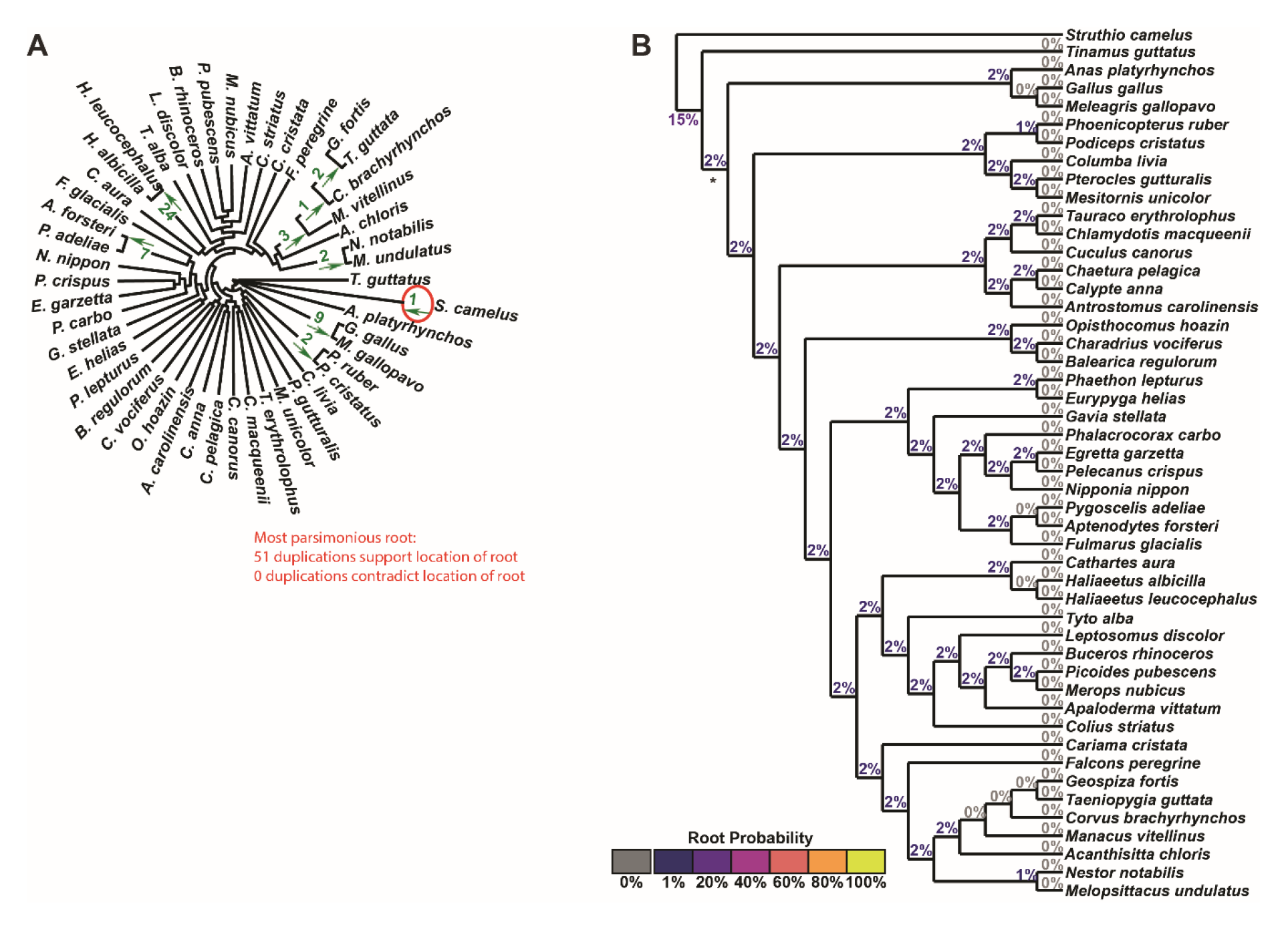
STRIDE analysis applied the set of Bird gene trees. A) Numbers of identified gene duplication events are marked on the branches they are observed on and arrows indicate the direction in which the duplication occurred. Gene duplication events are in agreement with the maximum parsimony root of the tree if the arrow points away from the root, and are in green. Those that disagree are in blue. The maximum parsimony root is circled in red and is in agreement with the correct root, marked with a *. B) The probabilities for the location of the root calculated by STRIDE, coloured according to the displayed heat map.

While STRIDE identified the community-accepted root in 75% of the datasets, it failed to identify this root for the bird (Fig. 7), rodent and Laurasiatheria (Supplementary File 1 Fig. S11 & S12) datasets. These three datasets had the smallest, second smallest and fourth smallest number of informative gene duplication events per species respectively (Table 1). In addition, while there were no conflicting gene duplication events in the bird dataset, the rodent and Laurasiatheria datasets had the highest and fifth highest ratio of conflicting to informative duplications (Table 1). Consistent with these observations, analysis of the factors affecting the accuracy of STRIDE revealed that root probability assignment was positively correlated with the number of informative duplications per species (R^2^=0.17, Supplementary File 1 Fig. S13A) and negatively correlated with the proportion of duplications which were in conflict (R^2^=0.24, Supplementary File 1 Fig. S13B). Furthermore, the proportion of conflicting duplications was negatively correlated with the number of species (R^2^=0.36, Supplementary File 1 Fig. S13C), suggesting increased taxon sampling facilitated more accurate identification of gene duplication events. Thus the ability of STRIDE to detect the true root is affected by taxon sampling and the number of gene duplication events detected in the dataset.

### STRIDE Provides Evidence for Location of the Root of the Eukaryotic Tree

Given the performance of stride on the datasets outlined above it was assessed whether STRIDE could provide insight into one of the most contentious and difficult tree rooting problems in biology, the root of the eukaryotic tree (Burki, F. 2014). Here, a set of 45 species that were distributed across the eukaryotic tree were selected. These were subject to orthogroup and gene tree inference as before and the complete set of 16770 gene trees were submitted for analysis by STRIDE. This identified 2316 putative gene duplication events excluding the root from (and supporting the monophyly of) major clades within the eukaryotes including the opisthokonta, fungi, metazoa, and achiplastida (Fig. 8A). Duplication events supporting further subclades within these major groupings were also abundant (Fig. 8A). In contrast, other major sub-clades including amoebazoa, the SAR supergroup, and the excavata, did not receive support from gene duplication events (Fig. 8A). This lack of gene duplication events meant that STRIDE could not exclude the root of the species tree from the basal branches of these groups and thus could not provide evidence for or against the five most popular placements for the root of the eukaryotic tree (Burki, F. 2014). This ambiguity in root assignment is represented effectively in the probabilities assigned to all putative root-spanning branches (Fig. 8B).

**Figure 8.**
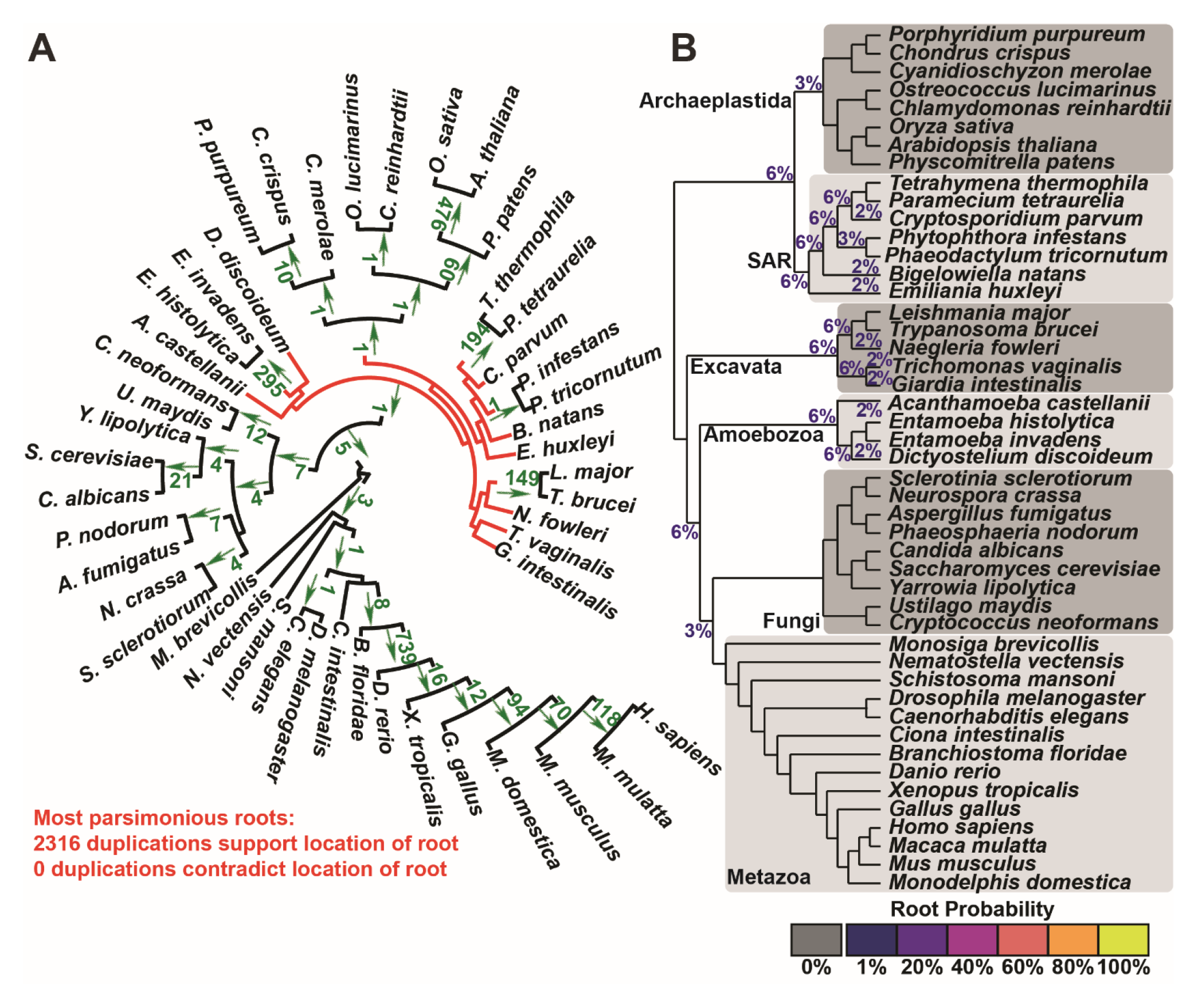
STRIDE analysis applied the set of Eukaryotic gene trees. A) Numbers of identified gene duplication events are marked on the branches they are observed on and arrows indicate which block of the bipartition the duplicate genes occur in. None of the gene duplication events contradict each other. The maximum parsimony roots have red branches, the branches from which the root is excluded are black. B) The probabilities for the location of the root calculated by STRIDE. Major groups of species are marked.

## Discussion

STRIDE is an automated method for identifying and analysing gene duplication events to infer the root of species trees. Through analysis of simulated and real datasets, we show how the performance of STRIDE is affected by data quantity, data conflict, and taxon sampling. Furthermore, we demonstrate that STRIDE is effective in identifying the root of species trees for the majority of species datasets and effectively captures the ambiguity in root assignment given the input data.

The aim of STRIDE is to infer a probability distribution over an entire species tree for the location of its root. This aim is different from algorithms that attempt to reconcile gene trees with species trees (Szollosi, G.J., Tannier, E., et al. 2015) or model duplication and loss processes on a tree (Gorecki, P. and Eulenstein, O. 2014). STRIDE identifies and utilises well-supported gene duplication events and does not evaluate gene loss events for the following reasons. First, gene trees can distinguish parallel duplication events on adjacent branches from a single shared duplication event, which is not possible for gene loss events. Second, the topology of the gene tree post-duplication genes can be compared with the species tree to confirm the accuracy of the inference, this cannot be done with gene loss events. Third, most genomes are incomplete and vary considerably in the quality of their annotation leading to high rates of false positive gene loss (Veeckman, E., Ruttink, T., et al. 2016, Dunne, M.P. and Kelly, S. 2017).

A major advantage of using STRIDE is that sets of species can be analysed without the inclusion on an outgroup. This is potentially advantageous in situations where inclusion of an outgroup can effect the topology of gene trees inferred for the ingroup species (Berger, S.A., Krompass, D., et al. 2011). Moreover, if the outgroup is distantly related to the ingroup species then additional problems of long branch attraction can lead to incorrect root placement (Philippe, H., Brinkmann, H., et al. 2011, Kuck, P., Mayer, C., et al. 2012, Salichos, L. and Rokas, A. 2013). STRIDE is also suitable for large dataset analysis and for situations where appropriate outgroups are not available. Although STRIDE as presented is a method for identifying the root of an unrooted species tree, the output from STRIDE can provide a wealth of useful information. For example, STRIDE maps high confidence gene duplication events to branches in a species tree. These gene duplication events provide strong evidence for monophyly of the species that share the gene duplication event. Thus STRIDE can be used to provide additional support to branches in a species tree that might be weakly supported by molecular sequence data. In this context, it is worth noting that STRIDE could also be used to evaluate support for alternative species-tree topologies by providing support for clades from gene duplication events.

The application of STRIDE to the eukaryotes was able to exclude the root of the eukaryotes from the opisthokonts and from a number of other groups, however STRIDE was unable to uniquely place the root as there were insufficient gene duplication events identified that could exclude the root from other portions of the tree. It is likely that poor taxon sampling for some of the groups (e.g. the amoebozoa and excavata), coupled with genome reduction associated with adaptation to parasitism in many of these species, impeded the discovery of these gene duplication events. With improved taxon sampling STRIDE may ultimately be able to provide further insight as to the location of the root of the eukaryotic tree. Furthermore, as STRIDE produces branch-level probabilities these could be combined with probabilities obtained from other analyses to perform a multi-data-type analysis of the origin of the eukaryotes.

In summary, STRIDE is a fast and effective method for genome scale phylogenetic analysis that can be used both to identify high confidence gene duplication events and identify the root of species trees without the requirement for an outgroup.

## References

Aslett M, Aurrecoechea C, Berriman M, Brestelli J, Brunk BP, Carrington M, Depledge DP, Fischer S, Gajria B, Gao X, et al. 2010. TriTrypDB: a functional genomic resource for the Trypanosomatidae. Nucleic Acids Res, 38:D457–D462.

Berger SA, Krompass D, Stamatakis A. 2011. Performance, Accuracy, and Web Server for Evolutionary Placement of Short Sequence Reads under Maximum Likelihood. Syst Biol, 60:291–302.

Boussau B, Szollosi GJ, Duret L, Gouy M, Tannier E, Daubin V. 2013. Genome-scale coestimation of species and gene trees. Genome Res, 23:323–330.

Burki F. 2014. The Eukaryotic Tree of Life from a Global Phylogenomic Perspective. Csh Perspect Biol, 6.

Dobzhansky T. 2013. Nothing in Biology Makes Sense Except in the Light of Evolution. Am Biol Teach, 75:87–91.

Donoghue MJ, Mathews S. 1998. Duplicate genes and the root of angiosperms, with an example using phytochrome sequences. Mol Phylogenet Evol, 9:489–500.

dos Reis M, Donoghue PCJ, Yang ZH. 2014. Neither phylogenomic nor palaeontological data support a Palaeogene origin of placental mammals. Biol Letters, 10.

Dunne MP, Kelly S. 2017. OrthoFiller: utilising data from multiple species to improve the completeness of genome annotations. bioRxiv.

Emms DM, Kelly S. 2015. OrthoFinder: solving fundamental biases in whole genome comparisons dramatically improves orthogroup inference accuracy. Genome Biol, 16.

Felsenstein J. 1981. Evolutionary Trees from DNA-Sequences - a Maximum-Likelihood Approach. J Mol Evol, 17:368–376.

Goodstein DM, Shu SQ, Howson R, Neupane R, Hayes RD, Fazo J, Mitros T, Dirks W, Hellsten U, Putnam N, et al. 2012. Phytozome: a comparative platform for green plant genomics. Nucleic Acids Res, 40:D1178–D1186.

Gorecki P, Eulenstein O. 2014. DrML: Probabilistic Modeling of Gene Duplications. J Comput Biol, 21:89–98.

Huelsenbeck JP, Bollback JP, Levine AM. 2002. Inferring the root of a phylogenetic tree. Syst Biol, 51:32–43.

Huerta-Cepas J, Serra F, Bork P. 2016. ETE 3: Reconstruction, Analysis, and Visualization of Phylogenomic Data. Mol Biol Evol, 33:1635–1638.

Huson DH, Scornavacca C. 2012. Dendroscope 3: An Interactive Tool for Rooted Phylogenetic Trees and Networks. Syst Biol, 61:1061–1067.

Jarvis ED, Mirarab S, Aberer AJ, Li B, Houde P, Li C, Ho SYW, Faircloth BC, Nabholz B, Howard JT, et al. 2014. Whole-genome analyses resolve early branches in the tree of life of modern birds. Science, 346:1320–1331.

Katoh K, Standley DM. 2013. MAFFT Multiple Sequence Alignment Software Version 7: Improvements in Performance and Usability. Mol Biol Evol, 30:772–780.

Kuck P, Mayer C, Wagele JW, Misof B. 2012. Long Branch Effects Distort Maximum Likelihood Phylogenies in Simulations Despite Selection of the Correct Model. Plos One, 7.

Larsson A. 2014. AliView: a fast and lightweight alignment viewer and editor for large datasets. Bioinformatics, 30:3276–3278.

Nguyen LT, Schmidt HA, von Haeseler A, Minh BQ. 2015. IQ-TREE: A Fast and Effective Stochastic Algorithm for Estimating Maximum-Likelihood Phylogenies. Mol Biol Evol, 32:268–274.

Parfrey LW, Lahr DJG, Knoll AH, Katz LA. 2011. Estimating the timing of early eukaryotic diversification with multigene molecular clocks. Proceedings of the National Academy of Sciences of the United States of America, 108:13624–13629.

Philippe H, Brinkmann H, Lavrov DV, Littlewood DTJ, Manuel M, Worheide G, Baurain D. 2011. Resolving Difficult Phylogenetic Questions: Why More Sequences Are Not Enough. Plos Biol, 9.

Salichos L, Rokas A. 2013. Inferring ancient divergences requires genes with strong phylogenetic signals. Nature, 497:327-+.

Simmons MP, Bailey CD, Nixon KC. 2000. Phylogeny reconstruction using duplicate genes. Mol Biol Evol, 17:469–473.

Sytsma KJ, Pires JC. 2001. Plant systematics in the next 50 years - re-mapping the new frontier. Taxon, 50:713–732.

Szollosi GJ, Tannier E, Daubin V, Boussau B. 2015. The Inference of Gene Trees with Species Trees. Syst Biol, 64:E42–E62.

Tange O. 2011. GNU Parallel - The Command-Line Power Tool. ;login: The USENIX Magazine, 36:42–47.

Veeckman E, Ruttink T, Vandepoele K. 2016. Are We There Yet? Reliably Estimating the Completeness of Plant Genome Sequences. Plant Cell, 28:1759–1768.

Williams TA, Heaps SE, Cherlin S, Nye TMW, Boys RJ, Embley TM. 2015. New substitution models for rooting phylogenetic trees. Philos T R Soc B, 370.

Wu YC, Rasmussen MD, Bansal MS, Kellis M. 2014. Most parsimonious reconciliation in the presence of gene duplication, loss, and deep coalescence using labeled coalescent trees. Genome Res, 24:475–486.

Yates A, Akanni W, Amode MR, Barrell D, Billis K, Carvalho-Silva D, Cummins C, Clapham P, Fitzgerald S, Gil L, et al. 2016. Ensembl 2016. Nucleic Acids Res, 44:D710–D716.

